# Functional investigation of conserved glutamate receptor subunits reveals a new mode of action of macrocyclic lactones in nematodes

**DOI:** 10.1101/2020.12.17.423223

**Authors:** Nicolas Lamassiaude, Elise Courtot, Angélique Corset, Claude L. Charvet, Cédric Neveu

**Author notes:** Co-corresponding authors (CLC), (CN).

## Abstract

Glutamate-gated chloride channels receptors (GluCls) are involved in the inhibition of neurotransmission in invertebrates and represent major molecular targets for therapeutic drugs. Among these drugs, macrocyclic lactones (MLs) are widely used as anthelmintic to treat parasitic nematodes impacting both human and animal health. Despite massive use of MLs since the 80’s, the exact molecular targets of these drugs are still unknown in many important parasite species. Among the GluCl subunit encoding genes, *avr-14*, *glc-2*, *glc-3* and *glc-4* are highly conserved throughout the nematode phylum. Using the *Xenopus* oocyte as an expression system, we pharmacologically characterized these GluCl subunits from the model nematode *Caenorhabditis elegans*, the human filarial nematode *Brugia malayi* and the horse parasitic nematode *Parascaris univalens.* In contrast with *C. elegans*, expression of parasitic nematode subunits as homomeric receptors was not reliable and needed glutamate application at the mM range to induce low currents at the nA range. However, the co-expression of GLC-2 and AVR-14B lead to the robust expression of ML-sensitive receptors for the three nematode species. In addition, we demonstrated that for *C. elegans* and *P. univalens,* GLC-2 co-assembled with GLC-3 to form a new GluCl subtype with distinct pharmacological properties. Whereas 1μM ivermectin, moxidectin and eprinomectin acted as agonist of the GLC-2/GLC-3 receptor from *C. elegans*, they did not directly activate GLC-2/GLC-3 of *P. univalens.* In contrast, these MLs potentialized glutamate elicited currents thus representing a unique pharmacological property. Our results highlight the importance of GLC-2 as a key subunit in the composition of heteromeric channels in nematodes and demonstrate that MLs act on novel GluCl subtypes that show unusual pharmacological properties, providing new insights about MLs mode of action.

**Author summary:** The filarial and ascarid parasitic nematodes include some of the most pathogenic or invalidating species in humans, livestock and companion animals. Whereas the control of these worms is critically dependent on macrocyclic lactones (MLs) such as ivermectin, the mode of action of this anthelmintic class remains largely unknown in these parasites. In the model nematode *Caenorhabditis elegans*, MLs target GluCl pentameric glutamate-sensitive chloride channels (GluCl). Because MLs are potent anthelmintics on *C. elegans*, ascarid and filarial nematodes, in the present study we investigated GluCl subunits highly conserved between these distantly related worms. Using the *Xenopus* oocyte as a heterologous expression system, we identified and performed the pharmacological characterization of novel GluCl receptors from *C. elegans*, the human filarial parasite *Brugia malayi* and the horse parasite *Parascaris univalens*. Our results highlight heteromeric GluCls from parasites as molecular targets for a wide range of MLs. We report an original mode of action of MLs on a new GluCl subtype made of the GLC-2/GLC-3 subunit combination. This study brings new insights about the diversity of GluCl subtypes in nematodes and opens the way for rational drug screening for the identification of next generation anthelmintic compounds.

## Introduction

The phylum *Nematoda* is divided into five major clades (I to V) that include free living and parasitic species impacting both human and animal health (1,2). Among these parasitic nematodes, *Filarioidea* and *Ascaridoidea* belonging to clade III are considered as the most impacting on both human and animal health (3). In this study, we focus our research on two parasitic nematode species representative of human filarids and animal ascarids: *Brugia malayi*, a human lymphatic filarid which is the causative agent of chronic elephantiasis in the south and south-east of Asia (4) and *Parascaris spp.* which are responsible for equine ascaridiosis (5,6).

Without effective vaccine or alternative strategies (7), the use of anthelmintic treatments remains the standard control strategy for parasitic nematodes. Among the available anthelmintic drugs, the broad-spectrum macrocyclic lactones (MLs) are highly effective and are massively used in human and veterinary medicine (8). The MLs include avermectins (ivermectin, doramectin, eprinomectin, abamectin, selamectin, emamectin) and milbemycins (moxidectin, milbemycin-oxime, nemadectin) that are potent against both endo- and ectoparasites (9). The morbidity and socio-economic impact of human lymphatic filariasis motivated control programs led by the World Health Organization (WHO) with ivermectin (IVM) as a spearhead of eradication operations (10). For the control of *Parascaris spp*. infestations, three drug classes currently have marketing approval including benzimidazoles, pyrantel and MLs (ivermectin and moxidectin), corresponding to the most widely used family. Unfortunately, because of the intensive use of MLs, treatment failures and resistant parasites have been reported worldwide (11). Resistance to MLs is spreading fast and has currently been reported in a wide range of parasitic nematode species such as *Onchocerca volvulus* (12), *Cooperia oncophora* (13,14), *Dirofilaria immitis* (15,16), *Haemonchus contortus* (17) and *Parascaris spp*. (18–25). Resistance is considered as multifactorial as it can raise from different molecular events such as: receptor subunit mutations (26–31), decrease of the target expression level (17,32) and the efflux mechanisms involving cell membrane transporter such as P-glycoprotein (33,34).

The molecular targets of the MLs as well as the mechanisms involved in resistance remains unclear for most of the parasitic nematodes. In this context, a better understanding of MLs mode of action is essential for the control of resistance and the development of novel therapeutical strategies (35).

In the free-living model nematode *Caenorhabditis elegans,* MLs act as allosteric modulators of glutamate-gated chloride channels (GluCls) (36). Exposure to MLs hyperpolarizes the membrane and inhibits the neurotransmission (37–39) leading to flaccid paralysis of the worms (26). GluCls are made of five subunits combining together to form either homo- or heteromeric receptors (40,41). In order to investigate the subunit composition and the pharmacological properties of recombinant nematode GluCls, the *Xenopus laevis* oocyte has proven to be an efficient heterologous expression system (42).

In *C. elegans*, six GluCl genes were identified and named *avr-14* (26,28), *avr-15* (38,43), *glc-1* (36,44), *glc-2* (36,43), *glc-3* (45) and *glc-4* (46). With the exception of GLC-4, all the subunits are able to form functional homomeric receptors when expressed in *Xenopus laevis* oocytes. All functional homomeric receptors were ivermectin-sensitive with the exception of Cel-GLC-2 which is not (36). However, it has been reported that the *C. elegans* GLC-2 subunit co-assemble with Cel-GLC-1 (36) or Cel-AVR-15 (43) to form two ivermectin-sensitive heteromeric GluCls subtypes with distinct pharmacological properties.

In contrast with *C. elegans*, only few functional GluCls have been characterized so far in parasitic nematodes. The AVR-14B subunit was reported to form a functional homomeric GluCls in *H. contortus* (28,47,48), *Cooperia oncophora* (27) and *Dirofilaria immitis* (49). Interestingly, in *Cooperia oncophora* (27) and *Haemonchus contortus* (50), GLC-2 also combined with AVR-14B to form a heteromeric GluCl subtype sensitive to ivermectin. Whereas GluCls investigations were mainly focused on clade V nematodes, the GluCl diversity and the mode of action of MLs remains poorly understood in human and animal parasitic nematodes from the clade III.

In the present study, we describe new GluCl subtypes made of highly conserved subunits from *C. elegans*, *B. malayi* and *P. univalens* providing new insights about the pharmacology of nematode GluCl subtypes as well as the mode of action of MLs.

## Results

### Four distinct GluCl subunits are conserved between *C. elegans*, *B. malayi* and *P. univalens*

Searches for homologs of GluCl subunits encoding genes from *C. elegans* (i.e. *glc-1*, *glc-2*, *glc-3*, *glc-4*, *avr-14* and *avr-15*) (26,36,38,43,45,46) in *B. malayi* and *P. univalens* genomic/transcriptomic databases, allowed the identification of 4 independent sequences corresponding to putative homologs of *avr-14*, *glc-2*, *glc-3* and *glc-4* from both species. In contrast, no homolog could be found in *B. malayi* and *P.* univalens for *glc-1* nor *avr-15*. The corresponding full-length cDNA of *avr-14*, *glc-2*, *glc-3* and *glc-4* from the three nematode species were cloned into a transcription vector for subsequent functional analysis. All GluCls identified including AVR-14B, GLC-2, GLC-3 and GLC-4 from *C. elegans*, *B. malayi* and *P. univalens* present the typical characteristics of a cys-loop receptor subunit. This includes, a predicted signal peptide in the extracellular N-terminal part (with the exception of Bma-GLC-2), the first cys-loop domains specific of ligand-gated ion channels (LGIC) which is composed of two cysteines that are 13 amino acid residues apart, the second cys-loop domain found in GluCls and four transmembrane domains (TM1-4) (**Fig. S2 and Fig. S3**).

In comparison with *C. elegans*, the GluCl deduced amino-acid sequences of *B. malayi* and *P. univalens* respectively show an identity of 78.4% and 80.3% for AVR-14B, 64.1% and 67.2% for GLC-2, 66.5% and 67.8% for GLC-3 and 64.8% and 66.6% for GLC-4 (**S1 Table**). A phylogenetic analysis including GluCl sequences from *C. elegans* (Cel), *B. malayi* (Bma), *H. contortus* (Hco) and *P. univalens* (Pun) confirmed the orthologous relationship of the parasitic subunit sequences with their respective counterparts in *C. elegans* (**Fig. 1)**. The identified sequences were then named according to the nomenclature proposed by Beech *et al* (51) and submitted to genbank under the accession numbers provided in the Material and Methods section.

**Fig. 1:**
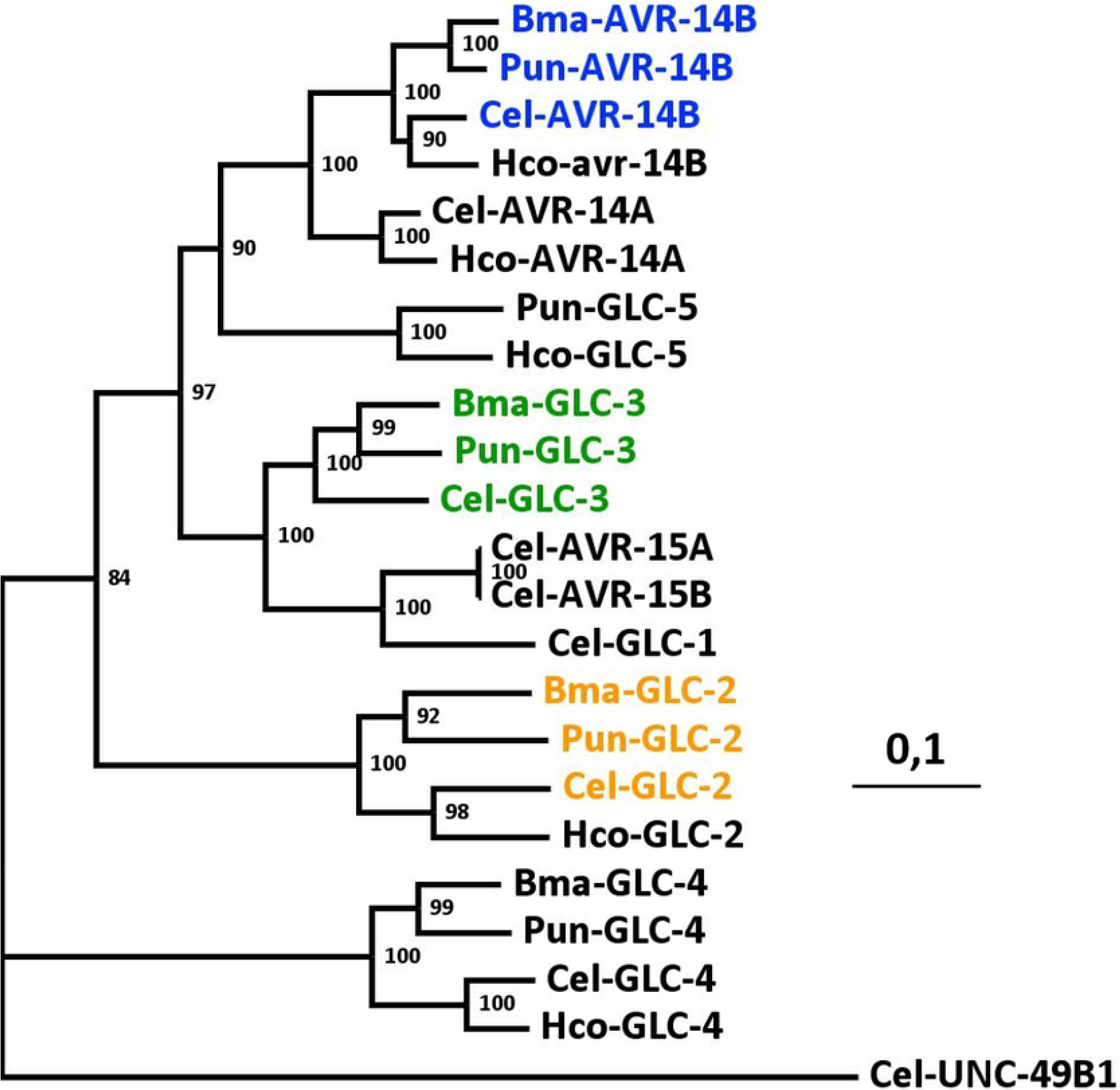
Distance tree of GluCl deduced amino-acid sequences from the nematodes *B. malayi* (Bma), *C. elegans* (Cel), *H. contortus* (Hco) and *P. univalens* (Pun). The bootstrap values (% from 1000 replicates) are indicated at each node. Scale bar represents the number of substitutions per site. Accession numbers for the sequences used in this analysis are provided in the Methods sections. Sequences of AVR-14B, GLC2 and GLC-3 from *B. malayi*, *C. elegans* and *P. univalens* are highlighted in blue, yellow and green respectively. The GABA receptor subunit UNC-49B from *C. elegans was* used as an outgroup.

### In contrast with *C. elegans*, GluCl subunits from *B. malayi* and *P. univalens* do not form robustly expressed homomeric receptors

In order to test the ability of *C. elegans*, *B. malayi* and *P. univalens* GluCl subunits to form functional homomeric receptors, their respective cRNA (*avr-14b, glc-2, glc-3* and *glc-4*) were injected singly in the *Xenopus laevis* oocyte. Three to five days post injection, currents elicited by 1 mM glutamate were recorded using the two-electrode voltage-clamp technic (**Fig. 2A**).

**Fig. 2:**
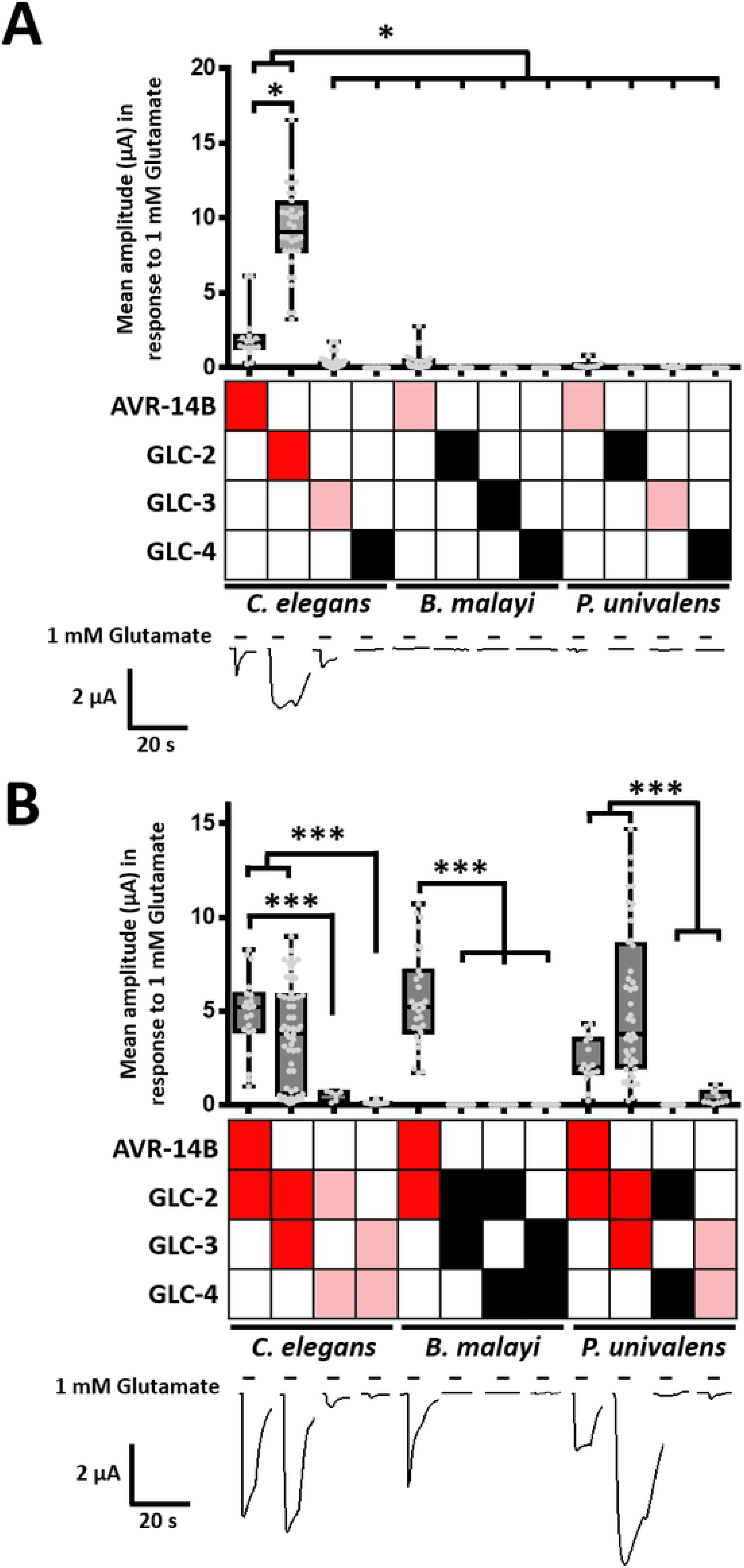
Functional expression of GluCls subunits from *C. elegans*, *B. malayi* and *P. univalens* in *Xenopus laevis* oocytes. Mean current amplitude in response to 1 mM glutamate application on *Xenopus* oocytes expressing homomeric (**A**) or heteromeric (**B**) receptors for *C. elegans*, *B.* malayi and *P. univalens.* Boxplots represent mean +/− SEM (*** p < 0.001). Dark red boxes show subunit combinations that led to robust expression of receptors responding to 1 mM glutamate with peak current in the μA range. Light red boxes correspond to combinations which respond to 1 mM glutamate with small currents in the nA range. Black boxes correspond to combinations that did not respond to 1 mM glutamate application. Representative glutamate-elicited currents traces are provided under each subunit combination, application times are indicated by the black bars.

Here, the *C. elegans* AVR-14B, GLC-2 and GLC-3 GluCl subunits were used as positive controls as their ability to form functional glutamate-sensitive homomeric channels has been previously reported (26,28,36,45). In our hands, expression of Cel-AVR-14B or Cel-GLC-2 alone formed functional homomeric receptors with robust glutamate-elicited currents in the μA range: 2.1 ± 0.4 μA (n = 16) and 9.1 ± 0.5 μA (n = 32) for Cel-AVR-14B and Cel-GLC-2 respectively (**Fig. 2A**). Whereas Cel-GLC-3 was also able to form functional receptors, the glutamate application (1 mM) induced significantly smaller peak currents (285 ± 54 nA, n = 50) in comparison with homomeric channels composed of Cel-AVR14B (p < 0.001) or Cel-GLC-2 (p < 0.001) (**Fig. 2A**). These results are in agreement with previous studies (26,36,45). Noteworthy, no glutamate-induced current was observed on oocytes injected with Cel-GLC-4 cRNA (n = 18).

Unlike *C. elegans,* where three out of four GluCl subunits formed functional homomeric receptors, for *B. malayi,* only the expression of AVR-14B subunit in oocytes allowed the reliable recording of currents (413 ± 132 nA, n = 23, **Fig. 2A**), significantly smaller than those obtained with Cel-AVR-14B (p < 0.001). Similarly, for *P. univalens*, the expression of AVR-14B subunit also allowed the formation of a homomeric receptor, with current amplitudes similar to the Bma-AVR-14B receptor (200 ± 80 nA, n = 9; p = 1, **Fig. 2A**). Furthermore, on oocytes expressing Pun-GLC-3, 1 mM glutamate application also resulted in low currents with mean amplitudes (77 ± 14 nA, n = 11) significantly smaller than for Cel-GLC-3 receptors (p < 0.05, **Fig. 2A**). Finally, in our hands, neither the expression of Bma-GLC-2 (n = 11), Pun-GLC-2 (n = 11), Bma-GLC-3 (n = 14), Bma-GLC-4 (n = 13) nor Pun-GLC-4 (n = 12) resulted in the expression of functional receptors.

Taken together, these results suggested that in contrast with *C. elegans*, parasitic nematode GluCl subunits are not able to form robustly expressed homomeric channels.

### GLC-2 has a critical role in heteromeric GluCl composition

Because previous studies highlighted the involvement of GLC-2 in heteromeric GluCls (27,36,43,50), oocytes expressing a combination of GLC-2 with either AVR-14B, GLC-3 or GLC-4 were challenged with 1 mM glutamate (**Fig. 2B**).

Firstly, we investigated the /AVR-14B/GLC-2 combination for the three different species. For *C. elegans*, the co-expression of both subunit cRNAs led to the robust expression of receptors with a mean current amplitude of 4.8 ± 0.4 μA (n = 21), significantly higher than Cel-AVR-14B alone (p < 0.05) but smaller than Cel-GLC-2 alone (p < 0.05). For *B. malayi*, the combination of AVR-14B and GLC-2 led to the robust expression of receptors sensitive to 1 mM glutamate (**Fig. 2B**) with a current peak of 5.6 ± 0.5 μA (n = 31), corresponding to current peaks 13-fold higher than for Bma-AVR-14B alone (p < 0.001). Similarly, for *P. univalens*, the combination of AVR-14B and GLC-2 led to the robust expression of functional receptors with 1 mM glutamate-elicited currents of 2.3 ± 0.3 μA (n = 17, **Fig. 2B**), 11-fold higher than for Pun-AVR-14B alone (p < 0.001).

These first results suggested that AVR-14B/GLC-2 were able to form heteromeric receptors potentially distinguishable from homomeric receptors.

Secondly, we tested the ability of the GLC-2 subunit to assemble with GLC-3. Strikingly, for *C. elegans,* we recorded strong currents elicited by 1 mM glutamate (3.7 ± 0.4 μA, n = 59) significantly different from homomeric receptors made of GLC-2 (p < 0.001) or GLC-3 (p < 0.001), thus suggesting that GLC-2/GLC-3 were able to form heteromeric receptors. Similarly, for *P. univalens,* combination of GLC-2/GLC-3 led to the robust expression of glutamate sensitive receptors with a current amplitude of 5.3 ± 0.6 μA (n = 45), significantly higher than the homomeric receptor formed by Pun-GLC-3 (p < 0.001, **Fig. 2B**). Noteworthy, Pun-GLC-2/GLC-3 receptor responses were similar to those of Cel-GLC-2/GLC-3 (p > 0.2). In contrast, for *B. malayi,* the GLC-2/GLC-3 combination failed to give rise to a functional receptor (n = 8).

Thirdly, we tested the combination of GLC-2 with GLC-4 for all species. Here, only very low currents (490 ± 106 nA, n = 14) were recorded from the Cel-GLC-2/GLC-4 combination (lower than for Cel-GLC-2 alone, p < 0.001) (**Fig. 2B**). Note that such a reduction of glutamate sensitivity has been previously reported, when GLC-2 was co-expressed with GLC-1 (36). In contrast with *C. elegans*, none of the Bma-GLC-2/GLC-4 and Pun-GLC-2/GLC-4 combination gave rise to glutamate-responsive receptor (n= 8 and n= 11 respectively).

Finally, we tested the combination of GLC-3 with GLC-4 subunit for all species. No currents were recorded with 1 mM glutamate application on oocytes injected with Bma-GLC-3/GLC-4 (n = 13), though nA currents were recorded for Cel-GLC-3/GLC-4 and for Pun-GLC-3/GLC-4. However, these currents were not different in comparison with homomeric GLC-3 channels (p = 1 and p = 0.12 for *C. elegans* and *P. univalens* respectively (**Fig. 2B**).

Taken together, these results demonstrate that GLC-2 from the three different species plays a pivotal role in the formation of glutamate-sensitive heteromeric receptor including AVR-14B or GLC-3.

### AVR-14B/GLC-2 from *C. elegans*, *B. malayi* and *P. univalens* form GluCl subtypes responsive to a wide range of macrocyclic lactones

As a first step, in order to distinguish the putative Cel-AVR-14B/GLC-2 heteromeric receptor from the homomeric Cel-AVR-14B and Cel-GLC-2, we challenged oocytes expressing both subunits singly or in combination and established their respective glutamate concentration-response curve. Representative currents induced by glutamate on Cel-AVR-14B/GLC-2 are shown in **Fig. 3A**. Glutamate EC_50_ values of 112.3 ± 13 μM (n = 5), 214.6 ± 13.1 μM (n = 5) and 20.5 ± 1.3 (n = 8) were determined for the Cel-AVR-14B, Cel-GLC-2 and Cel-AVR-14B/GLC-2 receptor respectively (**Fig. 3B**). Cel-AVR-14B/GLC-2 channel showed a higher sensitivity for glutamate in comparison with the homomeric receptors formed by AVR-14B (p < 0.001) or GLC-2 (p < 0.001), with 5- and 10-fold lower EC_50_, respectively. These results confirmed that Cel-AVR-14B/GLC-2 corresponds to a heteromeric GluCl receptor.

**Fig. 3:**
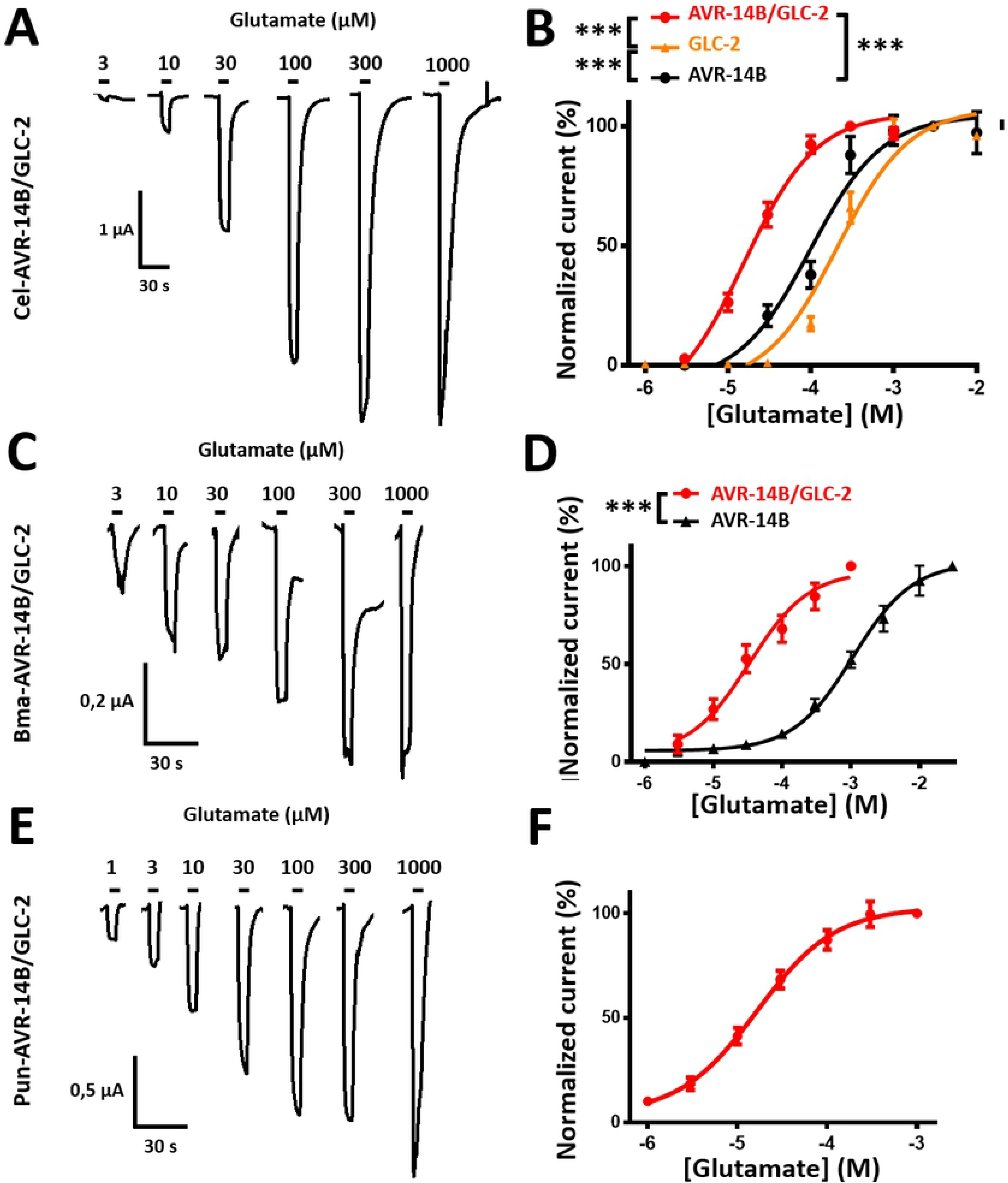
Functional characterization of AVR-14B/GLC-2 from *C. elegans, B. malayi and P. univalens*. **A.** Representative current traces of Cel-AVR-14B/GLC-2 expressed in *Xenopus* oocytes in response to the application of increasing concentrations of glutamate (3-1000 μM). Glutamate application times are indicated by the black bars. **B.** Glutamate concentration-response curve for Cel-AVR-14B, Cel-GLC-2 and the combination Cel-AVR-14B/GLC-2 (mean +/− SEM, n = 5-8). Current amplitudes were normalized to the maximal effect obtained with a saturating glutamate concentration (*** p < 0.001). **C.** Representative current traces of Bma-AVR-14B/GLC-2 expressed in *Xenopus* oocytes in response to the application of increasing concentrations of glutamate (3-1000 μM). Glutamate application times are indicated by the black bars. **D.** Glutamate concentration-response curve for Bma-AVR-14B and the combination Bma-AVR-14B/GLC-2 (mean +/− SEM, n = 5-7). Current amplitudes were normalized to the maximal effect obtained with a saturating glutamate concentration (*** p < 0.001). **E.** Representative current traces of Pun-AVR-14B/GLC-2 expressed in *Xenopus* oocytes in response to the application of increasing concentrations of glutamate (1-1000 μM). Glutamate application times are indicated by the black bars. **F.** Glutamate concentration-response curve for Pun-AVR-14B/GLC-2 (mean +/− SEM, n = 5). Current amplitudes were normalized to the maximal effect obtained with a saturating glutamate concentration.

For *B. malayi*, for which GLC-2 alone did not form a functional homomeric receptor, we determined the glutamate EC_50_ values for Bma-AVR-14B and Bma-AVR-14B/GLC-2 respectively. For the homomeric AVR-14B receptor glutamate EC_50_ was 758.9 ± 82 μM (n = 5), whereas for Bma-AVR-14B/GLC-2 glutamate EC_50_ was 32.5 ± 5.1 μM (n = 7), (**Fig. 2**; **Fig. 3C**). Such difference of glutamate sensitivity between these two receptors provides strong evidence that AVR-14B and GLC-2 subunits can associate to form a heteromeric GluCl subtype, distinct from the homomeric receptor made of AVR-14B (p < 0.001; **Fig. 3D**).

Strikingly, for the homomeric Pun-AVR-14B receptor, application of high concentrations of glutamate only induced small currents. Indeed, even with 100 mM glutamate applications (n = 7), the currents never reached a plateau value, suggesting that the receptor was not saturable. In sharp contrast, the co-injection of AVR-14B and GLC-2 led to the robust expression of a functional receptor sensitive to a micromolar range of glutamate with an EC_50_ value of 13.6 ± 1.6 μM (n = 5, **Fig. 3F**), thus confirming that Pun-GLC-2/AVR-14B form a heteromeric receptor (**Fig. 3E**).

Subsequently, in order to get new insight about the respective pharmacological properties of the heteromeric AVR-14B/GLC-2 GluCl subtypes from *C. elegans*, *B. malayi* and *P. univalens*, their sensitivity to a wide range of MLs available in the market has been investigated (i.e. abamectin, doramectin, emamectin, eprinomectin, ivermectin, moxidectin, and selamectin). Representative currents of the two most potent MLs on the AVR-14B/GLC-2 receptor from each species are presented in **Fig. 4A, Fig. 4C and Fig. 4E**. All the MLs tested acted as potent agonist on these receptors, inducing their permanent activation. Receptor activation by MLs was not reversible and slower in comparison with glutamate. Interestingly, whereas emamectin was found to be among the most potent ML on Cel-AVR-14B/GLC-2 and Bma-AVR-14B/GLC-2 (with 1 μM emamectin induced-current corresponding to 78 ± 7 % (n = 7) and 81 ± 6 % (n = 7) of the maximum current amplitude induced by glutamate, respectively), for Pun-AVR-14B/GLC-2 no significative difference of sensitivity was observed between the different MLs (**Fig. 4B, Fig. 4D and Fig. 4F**).

**Fig. 4:**
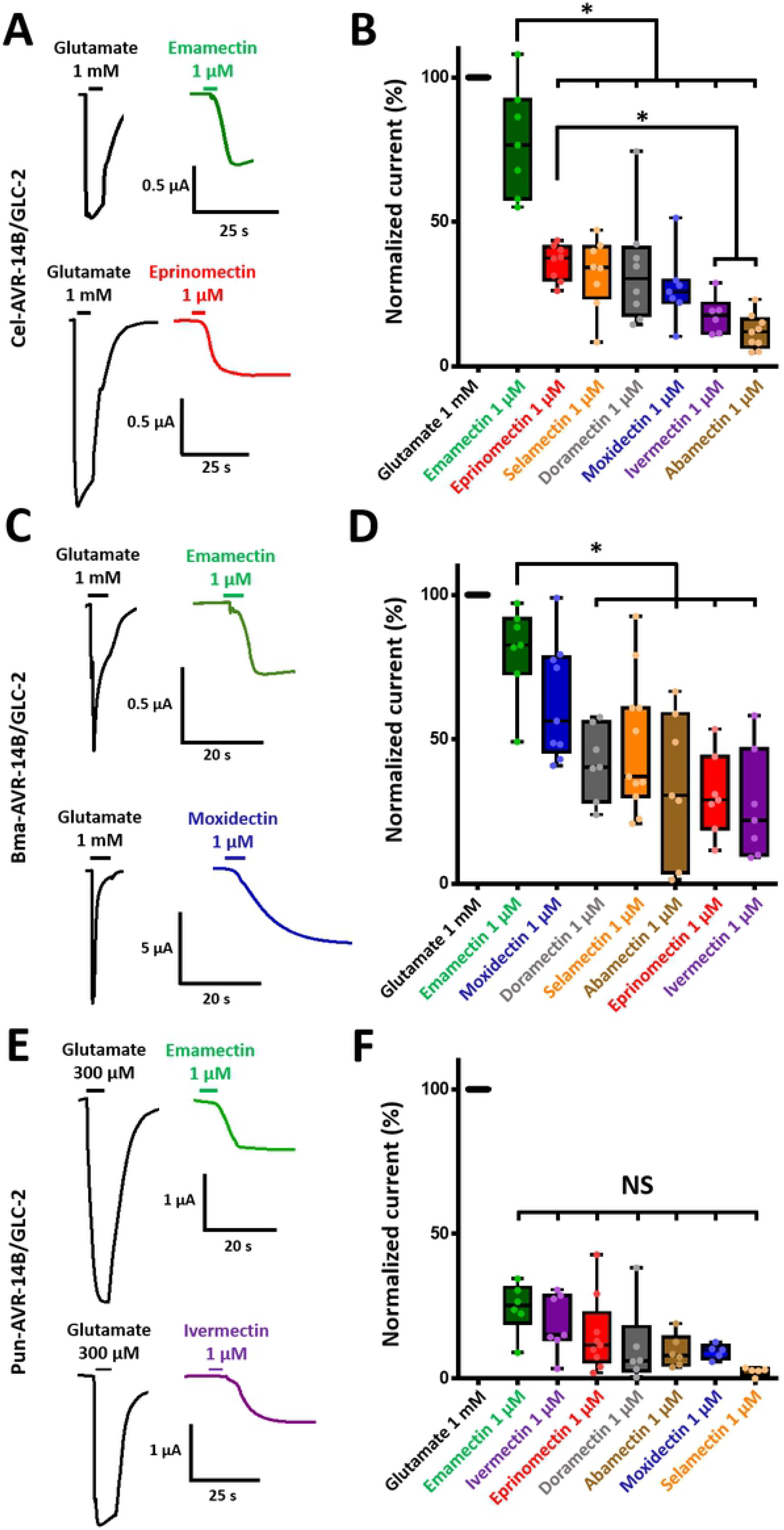
Effects of marketed macrocyclic lactones on AVR-14B/GLC-2 from *C. elegans*, *B. malayi* and *P. univalens*. **A.** Representative recording traces from a single oocyte injected with AVR-14B and GLC-2 of *C. elegans* induced by the two most potent activators, emamectin and eprinomectin after a first application of 1 mM glutamate. Application times are indicated by the black bars. **B.** Comparison of macrocyclic lactones effects at 1 μM after 5 s application on Cel-AVR-14B/GLC-2. All responses were normalized to the maximum responses obtained with glutamate at 1 mM (* p < 0.05). **C.** Representative current traces from a single oocyte injected with AVR-14B and GLC-2 of *B. malayi* induced by the two most potent activators, emamectin and moxidectin after a first application of 1 mM glutamate. Application times are indicated by the black bars. **D.** Comparison of macrocyclic lactones effects at 1 μM after 5 s application on Bma-AVR-14B/GLC-2. All responses were normalized to the maximum responses obtained with glutamate at 1 mM (* p < 0.05). **E.** Representative current traces from a single oocyte injected with AVR-14B and GLC-2 of *P. univalens* induced by the two most potent activators, emamectin and ivermectin after a first application of 1 mM glutamate. Application times are indicated by the black bars. **F.** Comparison of macrocyclic lactones effects at 1 μM after 5 s application on Pun-AVR-14B/GLC-2. All responses were normalized to the maximum responses obtained with glutamate at 1 mM (* p < 0.05).

### GLC-2/GLC-3 from *C. elegans* and *P. univalens* forms a new functional GluCl subtype with original pharmacological properties

As described for Cel-AVR-14B/GLC-2, in order to distinguish the putative Cel-GLC-2/GLC-3 heteromeric receptor from the homomeric Cel-GLC-2 and Cel-GLC-3 receptors, our objective was to determine and compare their respective glutamate EC_50_ values (**Fig. 5A**). For Cel-GLC-2/GLC-3, glutamate EC_50_ value was 46.8 ± 3.7 μM (n = 5), corresponding to a 4-fold reduction in comparison with the glutamate EC_50_ value previously determined for Cel-GLC-2 alone (214.6 ± 13.1 μM, n = 5). The shift between these two EC_50_ values clearly indicated that *C. elegans* GLC-2 and GLC-3 can combine to form a new GluCl subtype with a higher affinity for glutamate in comparison with the homomeric receptors (p < 0.001; **Fig. 5B**). Unfortunately, the maximum effect of glutamate application was not reached for Cel-GLC-3 (even with 30 mM glutamate), thus preventing the EC_50_ value determination for this receptor.

**Fig. 5:**
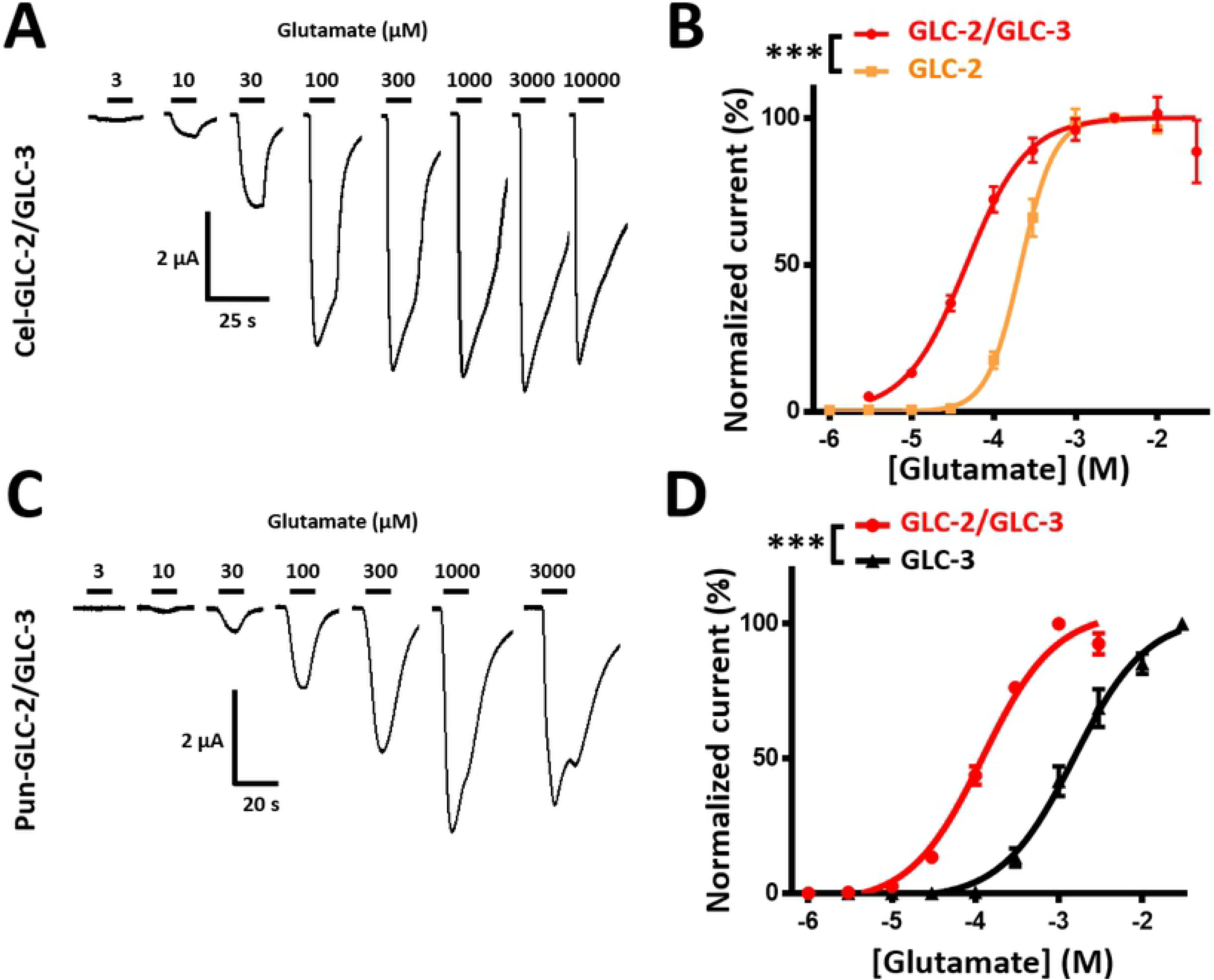
Pharmacological characterization of GLC-2/GLC-3 from *C. elegans* and *P. univalens*. **A.** Representative current traces of Cel-GLC-2/GLC-3 expressed in *Xenopus* oocytes in response to an increasing concentration of glutamate (3-10000 μM). Application times are indicated by the black bars. **B.** Glutamate concentration response curve on Cel-GLC-2 and on the combination Cel-GLC-2/GLC-3 (mean +/− SEM, n = 5). Current amplitudes were normalized to the maximal effect with glutamate (*** p < 0.001). **C.** Representative current traces of Pun-GLC-2/GLC-3 expressed in *Xenopus* oocytes in response to an increasing concentration of glutamate (3-3000 μM). Application times are indicated by the black bars. **D.** Glutamate concentration response curve on Pun-GLC-3 and on the combination Pun-GLC-2/GLC-3 (mean +/− SEM, n = 11-21). Current amplitudes were normalized to the maximal effect with glutamate (*** p < 0.001).

For *P. univalens*, the GLC-2/GLC-3 glutamate EC_50_ was 120.2 ± 5.7 μM (n = 21) whereas the EC_50_ for Pun-GLC-3 was 1482 ± 111 μM (n = 11) (**Fig. 5C, Fig.5D**). As mentioned for *C. elegans*, this drastic shift of EC_50_ values also confirmed that Pun-GLC-2/GLC-3 forms a novel subtype of heteromeric GluCl distinct from Pun-GLC-3 (p < 0.001).

In order to evaluate the involvement of GLC-2/GLC-3 as putative molecular targets for MLs, we decided to focused our attention on ivermectin, moxidectin and eprinomectin representing commonly used anthelmintic compounds (i.e. ivermectin and moxidectin used for *Parascaris spp.* treatment; eprinomectin used for clade V parasites on lactating animals).

The representative current traces obtained after drug applications on both Cel-GLC-2/GLC-3 and Pun-GLC-2/GLC-3 are shown in **Fig. 6A** and **Fig. 6C.** The Cel-GLC-2/GLC-3 receptor was activated by ivermectin, moxidectin and eprinomectin. The maximum current amplitude was induced by ivermectin with a current amplitude corresponding to 15 ± 2 % (n = of the maximum response obtained with glutamate (**Fig. 6B**) which is very similar to the ivermectin effect on Cel-AVR-14B/GLC-2 heteromeric receptor (18 ± 3 %, n = 6). Surprisingly, for *P. univalens*, none of the tested MLs induced a current (**Fig. 6D**). This result was further confirmed using a wider panel of MLs (**Fig. S4A**). In order to investigate if the drug application time could potentially impact the ML agonist effect, 1 μM ivermectin was perfused during 90 s on Pun-GLC-2/GLC-3. Even with this long-lasting application, ivermectin showed a weak agonist effect with a current amplitude corresponding to 3.56 ± 0.54 % (n = 7) of the maximum response obtained with glutamate (**Fig. S4B**).

**Fig. 6:**
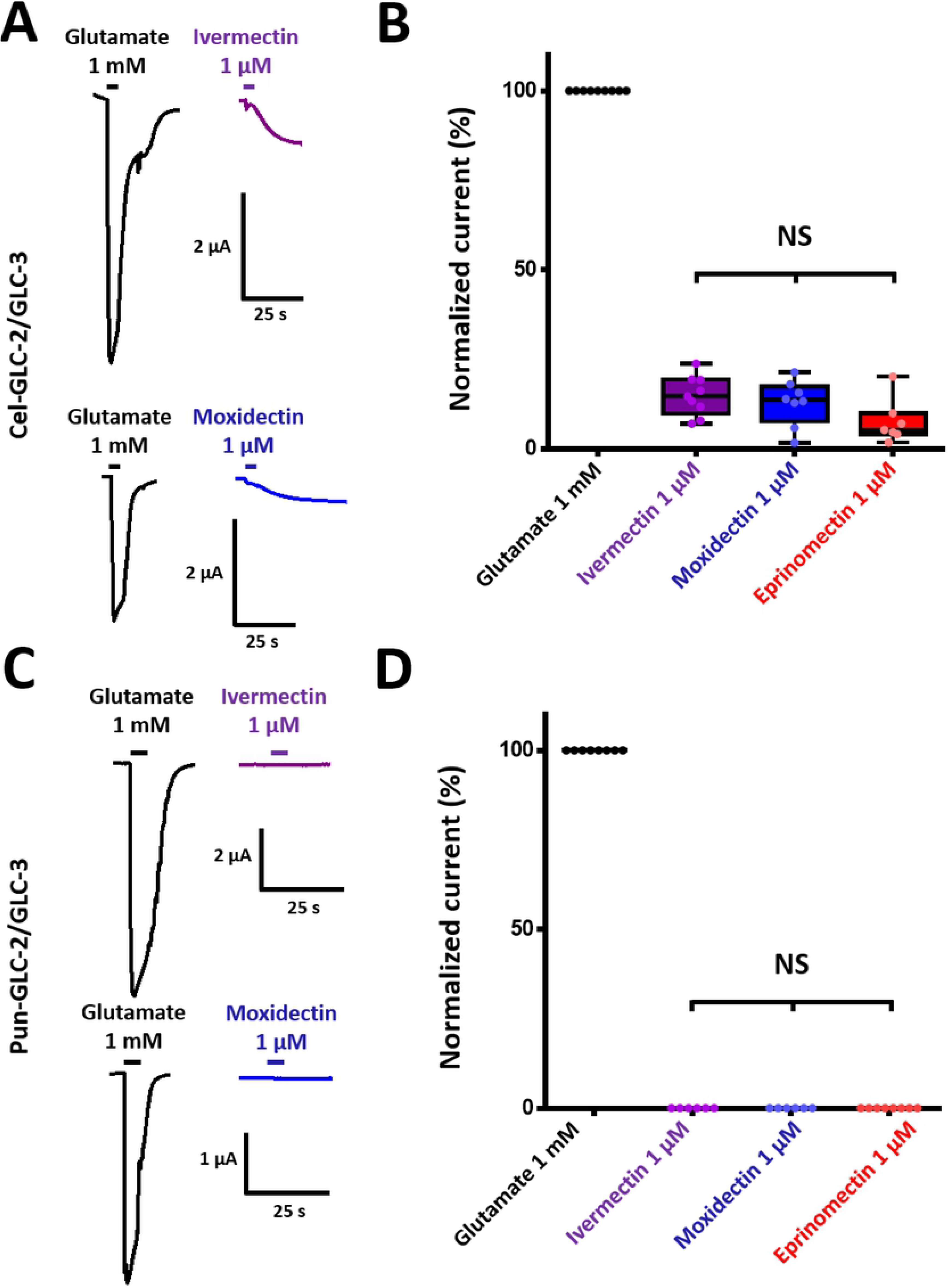
Pharmacological characterization of GLC-2/GLC-3 from *C. elegans* and *P. univalens*. **A.** Representative current traces induced by some macrocyclic lactones (ivermectin, moxidectin and eprinomectin) on Cel-GLC-2/GLC-3. Application time are indicated by the black bars. **B.** Comparison of some macrocyclic lactones effects at 1 μM after 5 s application on Cel-GLC-2/GLC-3. All responses are normalized to the maximum responses obtained with glutamate at 1 mM (NS = Not significative). **C.** Representative current traces induced by some macrocyclic lactones (ivermectin, moxidectin and eprinomectin) on Pun-GLC-2/GLC-3. Application time are indicated by the black bars. **D.** Comparison of some macrocyclic lactones effects at 1 μM after 5 s application on Pun-GLC-2/GLC-3. All responses are normalized to the maximum responses obtained with glutamate at 1 mM (NS = Not significative).

Because of this unexpected lack of activity as agonists, we hypothesized that MLs could potentially act as antagonists or potentialize the effect of glutamate on Pun-GLC-2/GLC-3. In order to test the hypothesis, 100 μM glutamate (corresponding approximately to the receptor glutamate EC_50_) were applied before, during and after the addition of 1 μM ivermectin (**Fig. 7A**) or moxidectin (**Fig. 7B**) on the receptor. Strikingly, both drugs potentialized the effect of glutamate on Pun-GLC-2/GLC-3. Current amplitudes in response to 100 μM glutamate were increased by 45 ± 10 % (n = 5, p < 0.05; **Fig. 7C**) and 93 ± 19 % (n = 8, p < 0.01; **Fig. 7D**) when co-applied with ivermectin or moxidectin respectively. This effect was reversible for both ivermectin (p < 0.05) and moxidectin (p < 0.001).

**Fig. 7:**
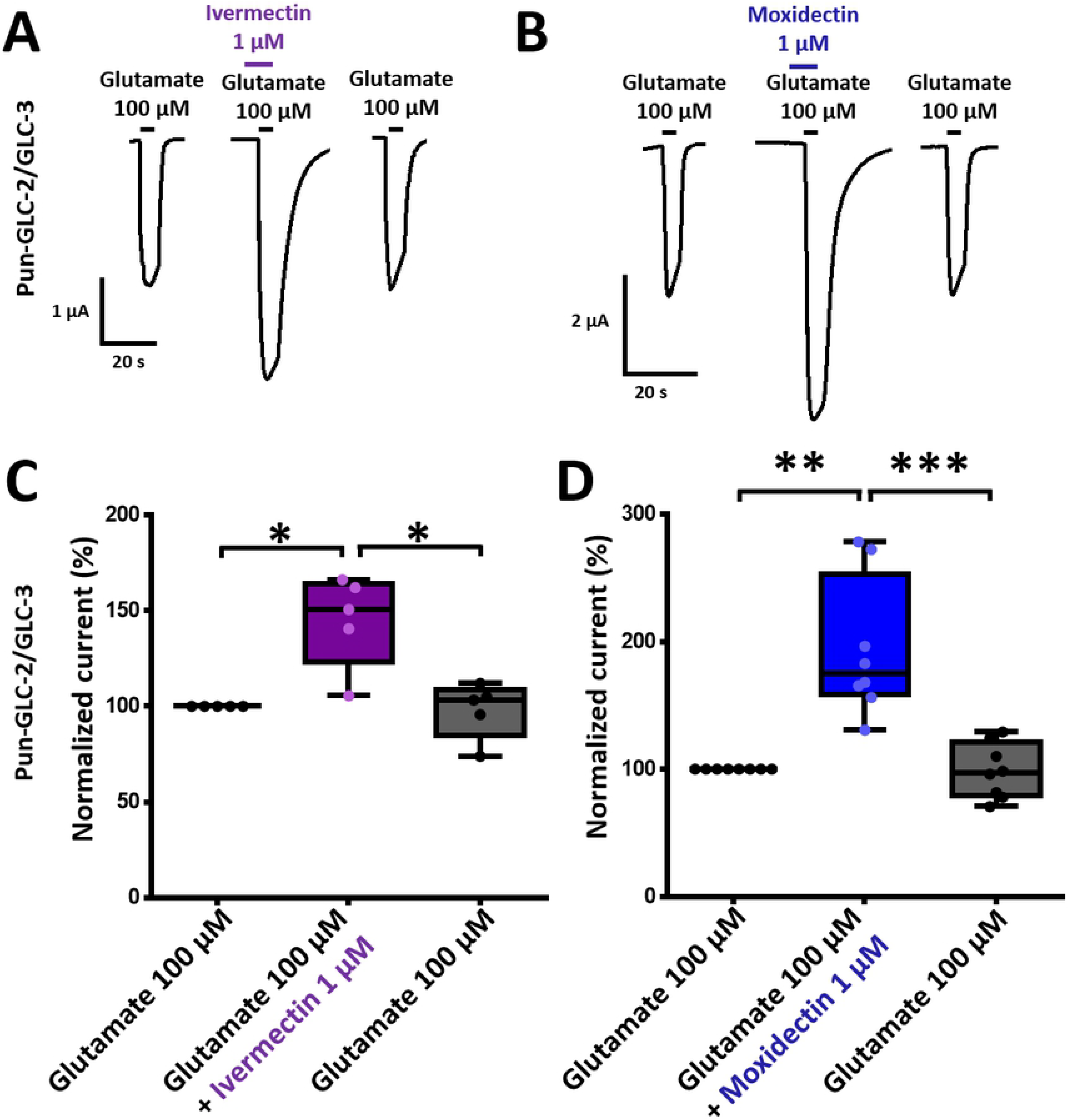
Modulation of glutamate effect by ivermectin and moxidectin on GLC-2/GLC-3 from P. univalens. **A-B.** Representative current traces induced by glutamate 100 μM followed by a co-application of ivermectin 1 μM (**A**) or moxidectin 1 μM (**B**) with glutamate 100 μM. Application times are indicated by the black bars. **C.** Box plot of the potentializing effect of ivermectin on Pun-GLC-2/GLC-3 normalized and compared with the response to 100 μM glutamate (* p < 0.05). **D.** Box plot of the potentializing effect of moxidectin on Pun-GLC-2/GLC-3 normalized and compared with the response to 100 μM glutamate (** p < 0.01, *** p < 0.001)

This is to our knowledge, the first report of a nematode GluCl potentialized by MLs at such concentration, representing a novel receptor subtype with unique pharmacological properties.

The pharmacological properties of the functional GluCl receptors described in the present work are summarized in S2 Table.

## Discussion

Among the six distinct genes encoding GluCl subunits *in C. elegans*, AVR-14, GLC-2, GLC-3 and GLC-4 are highly conserved in distantly related nematode species from different phylogenetic clades (52). Therefore, we reasoned that receptors including these subunits could be involved in the broad-spectrum activity of MLs on nematodes.

In the present study, using the xenopus oocyte as a heterologous expression system, we identified a panel of functional homomeric and heteromeric receptors made of the GLC-2, GLC-3 and AVR-14B subunits from the free-living model nematode *C. elegans* and *B. malayi* and *P. univalens*, two parasitic nematodes presenting a major impact for human and equine health respectively.

Among the *C. elegans* GluCl subunits, we confirmed that homomeric receptors made of AVR-14B or GLC-2 are responsive to glutamate in the μM range with elicited currents at the μA range, whereas in contrast, GLC-3 homomeric receptor only respond to mM glutamate application resulting in small currents at the nA range. Strikingly, for the latter, we showed that addition to GLC-2 led to the formation of a novel heteromeric GLC-2/GLC-3 receptor, responsive to more physiologically relevant glutamate concentrations. Even though it remains highly speculative to consider that such difference could reflect their existence *in vivo,* it clearly highlights the need to further investigate heteromeric GluCls as potential contributors to MLs sensitivity in nematodes.

In other clade V nematode species such as *H. contortus* (47,50) and *C. oncophora* (27) homomeric glutamate sensitive channels made of AVR-14B or GLC-2 have also been described, suggesting that such recombinant homomeric GluCl channels could also been obtained from other parasitic nematode species. However, in the present study, we showed that none of the GluCl subunits from the parasites *B. malayi* nor *P. univalens* (i.e. AVR-14B, GLC-2, GLC-3 and GLC-4) gave rise to robust functional channel when expressed in the *Xenopus* oocytes (i.e. no glutamate elicited current, or small current at the nA range elicited by mM range glutamate). These results further supported the need to explore heteromeric GluCls in target parasitic species.

### Highly conserved nematode GluCl subunits play a pivotal role in heteromeric receptors composition

In the present study, our hypothesis that GLC-2 could combine with other GluCls subunits was supported by previous studies reporting that GLC-2 can associate with GLC-1 or AVR-15 in *C. elegans* and with AVR-14B in *H. contortus* and *C. oncophora* (27,36,43,50). In addition, the *C. elegans* AVR-14, GLC-2, GLC-3 and GLC-4 subunits has been shown to be expressed in pharyngeal neurons, suggesting potential interactions between these subunits in the worm (53,54).

For *C. elegans*, *B. malayi* and *P. univalens*, the combination of AVR-14B/GLC-2 led to the robust expression of glutamate-sensitive receptors. The drastic reduction of the glutamate EC_50_ of AVR-14B/GLC-2 in comparison with their respective homomeric receptor counterparts strongly support the association of the two distinct subunits into functional heteromeric receptors. Subsequently, we described for the first time that GLC-2 and GLC-3 can associate to form a novel glutamate-sensitive GluCl subtype in *C. elegans* and *P. univalens*, but surprisingly not in *B. malayi*. Because Bma-GLC-2 has proven to be functional in the obligate heteromeric channel including AVR-14B, we first speculated that a putatively non-functional Bma-GLC-3 could be responsible for the failure to get the predicted GLC-2/GLC-3 receptor. However, the successfully expression of a functional chimeric receptor made of Pun-GLC-2 and Bma-GLC-3 (**Fig. S5**, n = 12), confirmed that Bma-GLC-3 was a functional subunit. Therefore, we could only speculate that additional subunits combining to GLC-3 alone or GLC-2/GLC-3 of *B. malayi* are required to form functional receptors. In addition, we cannot exclude that ancillary proteins might be required for the functional expression of GluCl receptor in *Xenopus* oocytes as reported for some acetylcholine-receptor subtypes (55,56).

In summary, our results highlight that subunit combination is critical for clade III parasites to form functional glutamate-sensitive receptor in *Xenopus* oocyte expression system but might also contribute to the diversity of GluCl subtypes in clade V nematodes.

### New insights about macrocyclic lactones mode of action

The identification of novel functional GluCl from three distinct nematode species opened the way for detailed pharmacological characterization to decipher their relative sensitivity to different MLs. In accordance with previous studies performed on AVR-14B/GLC-2 of *H. contortus* (50) and *C. oncophora* (27), we showed that AVR-14B/GLC-2 from *C. elegans*, *B. malayi* and *P. univalens* were sensitive to ivermectin and moxidectin corresponding to the two MLs with marketed authorization in human health (57) and the most widely used in equine health with abamectin and doramectin (58).

In addition, we showed that this GluCl subtype is also sensitive to a wide range of MLs including abamectin, doramectin, emamectin, eprinomectin and selamectin. Presently, only ivermectin has marketed authorization for a wide variety of hosts and parasites. Others MLs have more specific marketed authorization such as emamectin, which is mostly used as insecticide in veterinarian aquaculture as well as in terrestrial agriculture (59). Interestingly, the present work highlighted emamectin as the most efficient agonist of the AVR-14B/GLC-2 receptors in the three species in comparison with the currently used MLs against *B. malayi* and *Parascaris spp*. However, whether the high potency of emamectin on AVR-14B/GLC-2 could be correlated or not with an efficacy of this drug *in vivo* remains to be established. Importantly, this also raise the question of the relative contribution of the different nematode GluCl subtypes in MLs sensitivity. Indeed, in some case of ivermectin resistance, moxidectin remains effective to treat lambs infected with *H. contortus* (17) suggesting that both molecules could preferentially activate distinct pharmacological targets in the worms. Because stable transformation remains an elusive goal for numerous parasitic nematode species (60,61), RNAi experiments could represent an attractive alternative to investigate the respective role of the distinct GluCls subtype in MLs susceptibility (62). Recently RNAi has been successfully used in *B. malayi* to invalidate the expression of nAChR (63) and SLO-1 subunits (64). Undoubtedly, such an approach combined with phenotypic assays (65) or the recently developed *in vivo* imaging system (IVIS) optimized to study *B. malayi* on a gerbil model (66) would represent a major opportunity to investigate in more details the relative contribution of AVR-14B/GLC-2 in the MLs sensitivity.

### Distinct pharmacology between *C. elegans* and parasites

In the present study, we reported that GLC-2/GLC-3 of *C. elegans* and *P. univalens* form a new subtype of functional glutamate-sensitive receptors. However, depending on the nematode species, they presented very distinct pharmacological properties. Indeed, whereas ivermectin, moxidectin and eprinomectin act as agonist on Cel-GLC-2/GLC-3, in sharp contrast, these drugs had a reversible glutamate potentializing effect at the same concentration on the Pun-GLC-2/GLC-3 receptor. Such a pharmacological property appears to be rare in GluCls of invertebrates, opening the way for future investigations of GLC-2/GLC-3 in other parasitic species.

In conclusion, our study provides new insight about the GluCl diversity and highlights the importance of GLC-2 as a core subunit in heteromeric GluCls from *C. elegans* and the clade III nematodes *B. malayi* and *P. univalens*. This work opens the way for the systematic investigation of heteromeric GluCl subtypes in target parasitic species in order to lay a strong basis for the rational use of MLs and the discovery of novel drug targets for the development of next generation anthelmintics.

## Material and Methods

### A. Sample supply

*C. elegans* used in this study are Bristol N2 wild-type strain worms supplied by the *Caenorhabditis* Genetics Center (CGC), St. Paul, MN, USA, which is funded by NIH Office of Research Infrastructure Programs (P40 OD010440). *B. malayi* microfilariae were supplied by NIH/NIAID Filariasis Research Reagent Resource Center, University of Georgia, Athens, GA, USA (www.filariasiscenter.org). Adults *P. univalens* were collected in faeces of naturally infested foals from UEPAO (Experimental Unit of Orfrasière Animal Physiology, INRAE Centre Val de Loire, Nouzilly, 37380, France) 50 h after a treatment with ivermectin. All samples were store at −80°C in RNA later solution (Ambion) before used.

### B. Karyotyping of *Parascaris sp* samples

*Parascaris equorum* and *Parascaris univalens* are the two *Parascaris* species described as morphologically identical. They can be discriminated by karyotyping as *P. univalens* has one pair of chromosome while *P. equorum* has two pair (67). We confirmed the species status by karyotyping *Parascaris* eggs from foals in Nouzilly, France. Ascarid eggs were extracted from a pool of faecal samples from four foals from the UEPAO. Feces were mixed with tap water and deposited on two sieves stacking in order of size (125 μm on the top and 63 μm on the bottom, respectively). Eggs were collected and washed with a large amount of tap water on the 60 μm sieve. Karyotyping was performed as described previously (68). Briefly, eggs were decorticated by three washing steps with 2% sodium hypochlorite in 16.5% sodium chloride, and subsequently by six washing steps with cold tap water. Then, eggs were briefly centrifugated and split in pool of 1000-1500 eggs which were incubated for 1.5 h, 2 h or 3 h at 37 °C for the first or second embryonic division to occur. Eggs were fixed with a mixture of methanol, acetic acid and chloroform (6:3:1) during 1 h, then washed twice with tap water. Approximately 500 eggs were placed between a slide and a coverslip and were crushed by pressing hard manually 1 min on the slides and then frozen in liquid nitrogen for 1 min. The coverslip was removed and let the glass air dried. Dried slides and first coverslips were mounted with new coverslips and slides respectively, using ProLong® Diamond Antifade Mountant with DAPI (Life Technologies). Six mounted slides were incubated 24 h at room temperature in the dark and examined using a fluorescent microscope (Nikon Eclipse E600). As expected, the worldspread specie *P. univalens* was identified as the only specie present in the infected foals since all eggs had a single pair of chromosomes (**Fig. S1**).

### C. RNA extraction and cDNA synthesis

Total RNA was extracted from a pool of adults for *C. elegans* and from a pool of L4 larvae for *B. malayi*. For *P. univalens*, total RNA was extracted from the head of one worm, including pharynx. Total RNA was isolated with Trizol reagent (Invitrogen, Carlsbad, CA, USA) following the manufacturer’s recommendations. cDNA synthesis was performed with 0.5-5 μg of total RNA using the Maxima H minus Reverse Transcriptase kit (Thermo Scientific, Waltham, MA, USA) according to the manufacturer’s recommendations.

### D. Identification and cloning of full-length GluCls coding sequences from nematodes

PCR amplification were performed according to the manufacturer’s recommendations with the Phusion High Fidelity Polymerase (New England BioLabs, Ipswich, MA, USA) using cDNA as template. Full-length coding sequences were cloned into the transcription vector pTB-207 and RACE-PCR product were cloned into pGEM-T (Promega, Madison, WI, USA). Eurofins Genomics (Luxembourg, Luxembourg) sequenced all constructs. Sequences of Cel-AVR-14B (CAA04170), Cel-GLC-2 (AAA50786), Cel-GLC-3 (CAB51708) and Cel-GLC-4 (NP_495489.2) from *C. elegans* were available on Genbank as well as Pun-AVR-14B (ABK20343) subunit coding sequence from *P. univalens*. Using the GluCls sequences of *C. elegans*, *Haemonchus contortus* and Pun-AVR-14B as queries, tBLASTn searches in NCBI (http://blast.ncbi.nlm.nih.gov/Blast.cgi) and WormbaseParasite (https://parasite.wormbase.org/Tools/Blast?db=core) allowed the identification of full-length coding sequence of GluCls from *B. malayi* (Bma-GLC-2: XM_001893073.1; Bma-GLC-4: XM_001900205.1; Bma-AVR-14B: supercontig:Bmal-4.0:Bm_v4_Chr3_scaffold_001:1374074:1388079:1; Bma-GLC-3: Bmal-4.0:Bm_v4_Chr4_scaffold_001:1513094:1542759:-1) and partial sequences of GluCls from *P. univalens* (Pun-GLC-2: NODE_2545302; Pun-GLC-3: NODE_2129897 and NODE_2250308; Pun-GLC-4: NODE_1817943, NODE_2402242 and NODE_2418647). Primers designed for RACE PCR and for amplification of the full-length coding sequences of each subunit are indicated in **S3-5 Tables**. For GluCls subunits of *P. univalens*, the corresponding 5’ and 3’ cDNA ends were obtained by nested RACE-PCR experiments as previously described (56). After identification of the 5’ and 3’ ends, two pairs of new primers per subunit were designed to amplify the full-length coding sequence of all GluCl subunits by nested PCR with the proofreading Phusion High-Fidelity DNA Polymerase (Thermo Scientific). Then, PCR products were cloned into the transcription vector pTB-207 (69) using the In-Fusion HD Cloning kit (Clontech) as described previously (70). Recombinant constructs were purified using EZNA Plasmid DNA Mini kit (Omega Bio-Tek) and sequence-checked (Eurofins Genomics). The novel complete coding sequences of Bma-AVR-14B, Bma-GLC-2, Bma-GLC-3, Bma-GLC-4, Pun-AVR-14B, Pun-GLC-2, Pun-GLC-3 and Pun-GLC-4 were deposited to Genbank. Constructs were linearized with MlsI, PaeI or PacI restriction enzymes (Thermofisher) depending on the construct. Linearized plasmids were used as DNA templates for cRNA synthesis using the mMessage mMachine T7 transcription kit (Ambion). cRNAs were precipitated with lithium chloride and were resuspended in a suitable volume of RNAse-free water and stored at −20°C before use.

### E. Sequence analysis and phylogeny

Prediction of signal peptides and transmembrane domains were performed with SignalP4.0 (71) and Simple Modular Architecture Research Tool (72). Deduced amino-acid sequences of GluCls from *B. malayi*, *C. elegans*, *H. contortus* and *P. univalens* were aligned using the MUSCLE algorithm and further processed with GENEDOC (IUBio). Percentage of identity between deduced amino acid of mature protein without peptide signal were obtained with EBI Global Alignment EMBOSS Needle (73). The distance trees were constructed using the SeaView software (74) with BioNJ (Poisson) parameters. Significance of internal tree branches was estimated using bootstrap resampling of the dataset 1000 times. The tree was edited using the FigTree software (http://tree.bio.ed.ac.uk/software/figtree/).

The sequences used in this study are available on GenBank under the followed accession numbers: *Brugia malayi* (Bma): AVR-14B (MW196269), GLC-2 (MW196266), GLC-3 (MW196267), GLC-4 (MW196268); *Caenorhabditis elegans* (Cel): AVR-14A (AAC25481), AVR-14B (MW196270), AVR-15A (CAA04171), AVR-15B (CAA04170), GLC-1 (AAA50785), GLC-2 (AAA50786), GLC-3 (CAB51708), GLC-4 (NP_495489.2), UNC-49B1 (AAD42383); *Haemonchus contortus* (Hco): AVR-14A (CAA74622), AVR-14B (CAA74623), GLC-2 (CAA70929), GLC-4 (ABV68894), GLC-5 (AAD13405); *Parascaris univalens* (Pun): AVR-14B (MW187941), GLC-2 (MW187938), GLC-3 (MW187939), GLC-4 (MW187940) and GLC-5 (QBZ81966).

### F. Electrophysiological recording and data analysis in *Xenopus laevis* oocytes

Defolliculated *Xenopus laevis* oocytes were purchased from Ecocyte Bioscience (Germany) and maintained in incubation solution (100 mM NaCl, 2 mM KCl, 1.8 mM CaCl_2_.2H_2_O, 1 mM MgCl_2_.6H_2_O, 5 mM HEPES, 2.5 mM C_3_H_3_NaO_3_, pH 7.5 supplemented with 100 U/mL penicillin and 100 μg/mL streptomycin) at 19 °C. Using the Drummond nanoject II microinjector, each oocyte was microinjected with 72 ng of cRNA when subunits were expressed singly or with 50 ng for each subunit when expressed in combination (1:1 ratio). Three to six days after cRNA microinjection, two-electrode voltage-clamp recordings were performed with an Oocyte clamp OC-725C amplifier (Warner instrument) at a holding potential of −80 mV to assess the expression of the GluCl channels. Currents were recorded and analyzed using the pCLAMP 10.4 package (Molecular Devices).

Dose responses relationships for glutamate were carried out by challenging oocytes with 5-10 s applications of increasing concentration of glutamate (between 1 μM to 30 mM depending on the receptor) with 2 min washing steps with Ringer solution between each application. The peak current values were normalized to the maximum response obtained with a saturated concentration of glutamate. The concentration of agonist required to mediate 50% of the maximum response (EC_50_) and the Hill coefficient (nH) were determined and compared using non-linear regression on normalized data with R Studio using the *drc* package v3.0-1 (75). Results are shown as mean ± SEM.

Comparison of the MLs effect on a receptors were performed as described previously (27,49). Briefly, 1 mM glutamate was first perfused on the oocytes as a reference peak current (maximum response) before application of a ML at 1 μM for 5 s. Current responses were normalized to the maximum current amplitude obtained with 1 mM glutamate. For Pun-GLC-2/GLC-3, the potentializing effect of MLs was evaluated by a first application of each MLs alone for 5 s, followed by the co-application with 100 μM glutamate for 5 s. The observed responses were normalized to the response induced by 100 μM glutamate (corresponding approximately to the glutamate EC_50_ for Pun-GLC-2/GLC-3) alone performed prior to challenging with the ML. In order to investigate the reversibility of the potentializing effect of the MLs, 100 μM glutamate was applied after 2 min washing. Statistical analyses were performed using Wilcoxon’s test with Bonferroni adjustments to compared glutamate and MLs current amplitudes.

### G. Chemicals

Glutamate, piperazine and the macrocyclic lactones (selamectin, ivermectin, doramectin, emamectin, eprinomectin, abamectin and moxidectin) were purchased from Sigma-Aldrich. Macrocyclic lactones were first dissolved in DMSO as 10 mM and then diluted in recording solution to the required concentration with a final concentration of DMSO which not exceed 1 %. Glutamate was directly prepared in recording solution.

## Acknowledgments

We thank Pr. Adrian Wolstenholme from the Department of Infectious Diseases, College of Veterinary Medicine, University of Georgia, Athens, GA, 30602, USA for providing the *B.malayi* worms and the careful and critical reading of the manuscript. We thank the group of UEPAO (Experimental Unit of Orfrasière Animal Physiology, INRAE Centre Val de Loire, Nouzilly, 37380, France) for providing *P. univalens* eggs and adults.

## Supporting information

**Fig. S1: Karyotype of *Parascaris univalens***

*Parascaris* eggs were DAPI-stained during the first mitotic division. The single pair of chromosomes is representative of *P.univalens* (x400).

**Fig. S2: Amino-acid alignments of AVR-14B (A), GLC-2 (B) and GLC-3 (C) subunit sequences from the four nematode species *Brugia malayi* (Bma), *Caenorhabditis elegans* (Cel), *Haemonchus contortus* (Hco) and *Parascaris univalens* (Pun)**

Predicted signal peptides in the N-terminal region are highlighted in grey. Amino acids share between the four species are highlight in blue. The four transmembrane segments (TM 1-4) and cys-loops are indicated by the black bars.

**Fig. S3: Amino-acid alignment of GLC-4 from the four nematode species *Brugia malayi* (Bma), *Caenorhabditis elegans* (Cel), *Haemonchus contortus* (Hco) and *Parascaris univalens* (Pun)**

Predicted signal peptides in the N-terminal region are highlight in grey. Amino acids share between the four species are highlight in blue. The four transmembrane segments (TM 1-4) and cys-loops are indicated by the black bars.

**Fig. S4: Pharmacological characterization of GLC-2/GLC-3 from *P. univalens***

**A.** Comparison of the macrocyclic lactones effects at 1 μM after 5 s application on Pun-GLC-2/GLC-3. All responses are normalized to the maximum responses obtained with glutamate at 1 mM.

**B.** Representative current traces induced by ivermectin at 1 μM during 90 s on Pun-GLC-2/GLC-3. Application time is indicated by the black bar.

**Fig. S5: Representative current traces after application of glutamate 1 mM on *Xenopus* oocytes expressing chimera receptors made of Pun-GLC-2 and Bma-GLC-3 (n = 12)**

Application time are indicated by the black bars.

**S1 Table: Percentage amino acid sequence identity between GluCl subunits from *C. elegans*, *B. malayi* and *P. univalens***

**S2 Table: Effect of glutamate and macrocyclic lactones on GluCls from *C. elegans*, *B. malayi* and *P. univalens* expressed in *Xenopus* oocytes.**

**S3 Table: List of primers used for PCR and cloning experiments of GluCls subunits from *elegans*.**

**S4 Table: List of primers used for PCR and cloning experiments of GluCls subunits from *Brugia malayi***.

**S5 Table: List of primers used for PCR and cloning experiments of GluCls subunits from *Parascaris univalens***.

